# A *Drosophila* model to screen Alport syndrome *COL4A5* variants for their functional pathogenicity

**DOI:** 10.1101/2024.03.06.583697

**Authors:** Jianli Duan, Pei Wen, Yunpo Zhao, Joyce van de Leemput, Jennifer Lai Yee, Damian Fermin, Bradley A Warady, Susan L Furth, Derek K Ng, Matthew G Sampson, Zhe Han

## Abstract

Alport syndrome is a hereditary chronic kidney disease, attributed to rare pathogenic variants in either of three collagen genes (*COL4A3/4/5*) with most localized in *COL4A5*. Trimeric type IV Collagen α3α4α5 is essential for the glomerular basement membrane that forms the kidney filtration barrier. A means to functionally assess the many candidate variants and determine pathogenicity is urgently needed. We used *Drosophila*, an established model for kidney disease, and identify *Col4a1* as the functional homolog of human *COL4A5* in the fly nephrocyte (equivalent of human podocyte). Fly nephrocytes deficient for *Col4a1* showed an irregular and thickened basement membrane and significantly reduced nephrocyte filtration function. This phenotype was restored by expressing human reference (wildtype) *COL4A5*, but not by *COL4A5* carrying any of three established pathogenic patient-derived variants. We then screened seven additional patient *COL4A5* variants; their ClinVar classification was either likely pathogenic or of uncertain significance. The findings support pathogenicity for four of these variants; the three others were found benign. Thus, demonstrating the effectiveness of this *Drosophila* in vivo kidney platform in providing the urgently needed variant-level functional validation.

**SUMMARY STATEMENT:** *Drosophila*, an established model of kidney disease, was used to develop an in vivo functional screen to determine causation for *COL4A5* genetic variants linked to Alport syndrome, a progressive nephropathy.

## INTRODUCTION

Alport syndrome, also known as hereditary nephritis, is a rare progressive kidney disease characterized by hematuria and proteinuria that often presents with hearing loss and ocular abnormalities (Alport, 1927; Flinter, 1997; Hertz et al., 2015; Spear and Slusser, 1972). Kidney pathology is marked by glomerular basement membrane (GBM) splitting and lamellation (Barsotti et al., 2001; Kalluri et al., 1997; Longo et al., 2006; Noël, 2000). The GBM lies between the capillary epithelium and the podocyte foot processes and is an essential part of the kidney filtration unit. Key components of its scaffolding are Collagen IV α3α4α5 trimers (Naylor et al., 2021). Alport syndrome is caused by mutations in the genes that encode these Collagen type IV alpha proteins (*COL4A3*, *COL4A4*, and *COL4A5*) (Artuso et al., 2012; Barker et al., 1990; Cameron-Christie et al., 2019; Fallerini et al., 2014; Hadjipanagi et al., 2022; Heiskari et al., 1996; Hudson, 2004; Longo et al., 2006; Pokidysheva et al., 2021; Zhang et al., 2019). The mutations inhibit trimeric protein complex formation which prevents a pivotal developmental switch from Collagen type IV α1 and α2 isoforms in fetal kidney to the α3, α4, and α5 isoforms in mature podocytes (Harvey et al., 1998; Kalluri et al., 1997; Miner and Sanes, 1994). This leaves the collagen scaffold more vulnerable to proteolysis by collagenases and cathepsins, which are required for GBM maintenance and turnover during normal conditions (Gunwar et al., 1998; Zeisberg et al., 2006). Over time, the GBM deteriorates, resulting in the characteristic GBM splitting and lamellation observed in Alport patient kidney biopsies (Barsotti et al., 2001; Kalluri et al., 1997; Longo et al., 2006; Noël, 2000). To date, nearly 2,000 variants in the *COL4A(3,4,5)* genes have been linked to Alport syndrome: Most mutations are in the X-linked *COL4A5* gene (Daga et al., 2022; Savige et al., 2021), mutations in *COL4A3* and *COL4A4* on chromosome 2 are often autosomal recessive (Daga et al., 2022); in addition, oligogenicity has been reported (Daga et al., 2022; Savige et al., 2021; Zhang et al., 2021).

One of the biggest challenges facing nephrologists today is determining whether variants of unknown significance in *COL4A* genes found in patients with glomerulonephritis or proteinuric kidney diseases are actually causing/contributing to the patient’s condition. A reliance on bioinformatic predictions to determine pathogenicity is imperfectly accurate, which is not acceptable when making patient-care decisions. The availability of variant-specific functional data would address this need. Nephrocytes are the *Drosophila* equivalent of mammalian podocytes as both have dynamic slit diaphragm structures that carry out the critical filtration functions to maintain water and electrolyte homeostasis in the blood (Weavers et al., 2009). Even though the fly nephrocyte effectively has an inside-out filtration structure, consisting of lacuna channels and a basement membrane, nephrocytes and podocytes share many molecular and ultrastructural features. In fact, most genes associated with kidney disease in patients have functional homologs in the fly nephrocyte (Fu et al., 2017; Rani and Gautam, 2018) and in both kidney cells endocytosis and exocytosis are essential for the formation and maintenance of the slit diaphragm filtration structure (Lang et al., 2022; Wang et al., 2021; Weavers et al., 2009; Zhuang et al., 2009). The fly system has already shown its effectiveness in an in vivo functional renal gene discovery screen (Fu et al., 2017; Hermle et al., 2017; Rani and Gautam, 2018; Zhang et al., 2013). Here, we use *Drosophila* to develop an efficient screening platform to provide functional validation for patient derived *COL4A5* variants associated with Alport syndrome using data from participants in the Chronic Kidney Disease in Children (CKiD) cohort.

## RESULTS

### *Drosophila Col4a1* deficiency results in dysfunctional nephrocytes

Human *COL4A5* encodes the collagen type IV alpha 5 chain (COL4A5) protein. *Drosophila* Col4a1 shares the main protein features with human COL4A5, which includes an N-terminus signal peptide, and the characteristic long triple-helical collagenous domain which is flanked by the short N-terminal 7S domain, and the duplicated non-collagenous domain (NC1, a.k.a. C4) at the C-terminus (Figure 1A). We used the *Klf15*-Gal4 driver to knock down *Col4a1* by expressing RNAi (*Col4a1*-IR). We assayed nephrocyte uptake function using 10kD Dextran particles, and the much larger FITC-albumin (66kD), which is among the largest particles that can cross the slit diaphragm. Both *Col4a1*-IR lines tested, each carrying an independent hairpin design, showed significantly reduced uptake of 10kD Dextran particles (Figure 1B,C) and FITC-albumin (Figure 1D,E). Similar, the uptake of the *Mhc*-ANF-RFP reporter (*Myosin heavy chain* promoter region drives the expression of full-length Rnor\Nppa cDNA, tagged with DsRed(T4) fluorescent protein) was significantly decreased in nephrocytes deficient for *Col4a1* (Supplemental Figure 1). These results show that, like its homolog COL4A5 in human podocytes, Col4a1 is crucial for nephrocyte function in flies.

**Figure 1.**
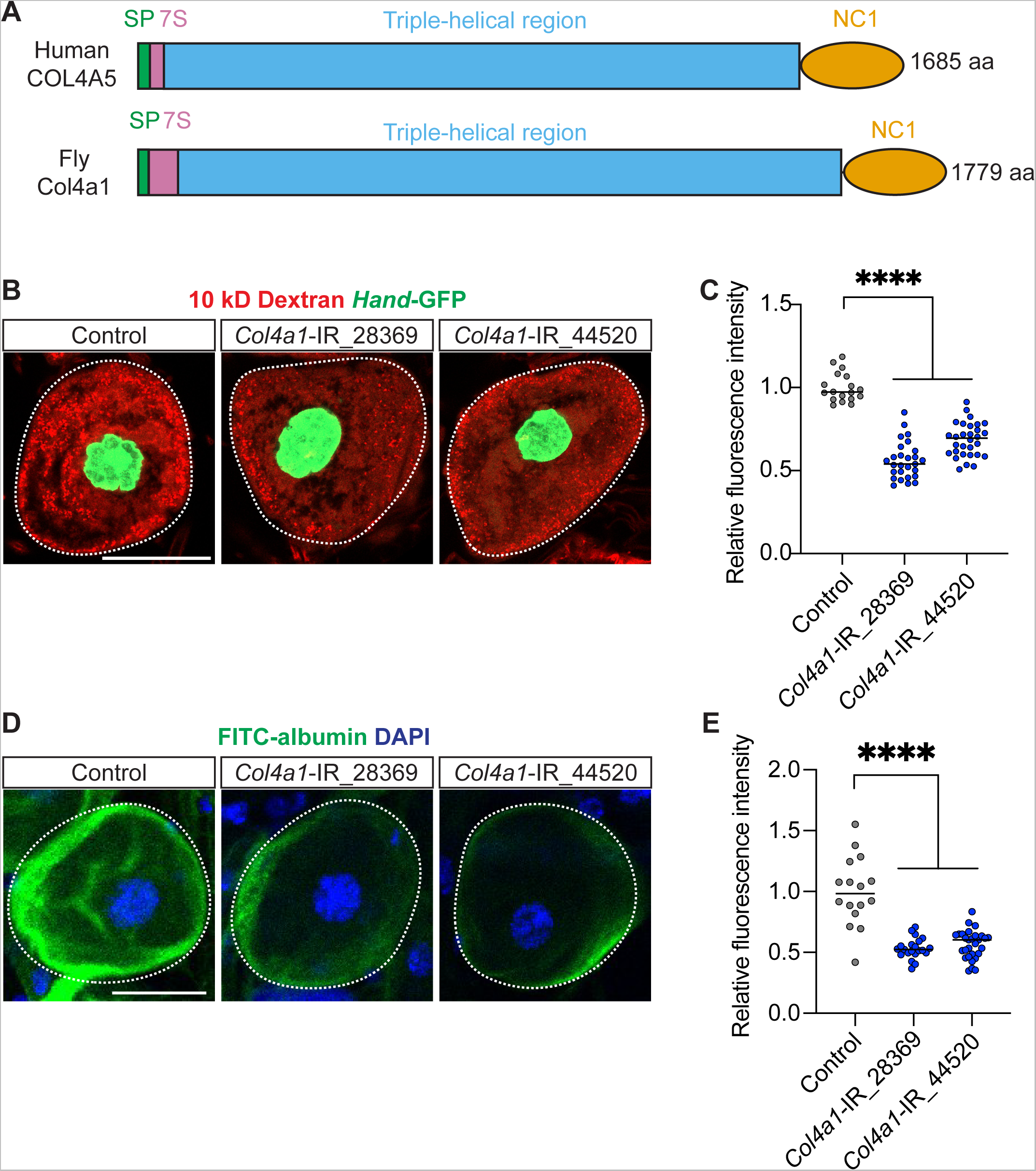
*Col4a1*-deficiency in fly nephrocytes causes functional defects. (**A**) Graphic display of protein domains for human COL4A5 and fly Col4a1: signal peptide (SP), seven Svedberg units (7S), triple helical region, and non-collagenous (NC1) domains. aa, amino acids. (**B**) Representative confocal images of 10 kD Dextran uptake (red) by female adult nephrocytes from Control and *Col4a1*-IR fly lines. *Hand*-GFP transgene expression was visualized as green fluorescence concentrated in the nephrocyte nuclei. Scale bar: 20 µm. (**C**) Quantitation of the relative fluorescence intensity of 10 kD Dextran in (B). (**D**) Representative confocal images of FITC-albumin uptake by female adult nephrocytes from Control and *Col4a1*-IR fly lines. DAPI (blue) indicates nuclei. Scale bar: 20 μm. (**E**) Quantitation of the relative fluorescence intensity of FITC-albumin in (D). (**B-E**) Flies: Control, (*Hand*-GFP/+; *Klf15*-Gal4/+); and *Col4a1*-IR (VDRC_28369 or BDSC_44502), (*Hand*-GFP/+; *Klf15*-Gal4/+; UAS-*Col4a1*-IR/+). (**C,E**) Statistical significance was defined as P<0.05 using one-way ANOVA with Tukey multiple comparisons test; (****) signifies P<0.0001.

### *Drosophila* Col4a1 is the functional homolog of human COL4A5 in nephrocytes

To verify that fly Col4a1 is indeed the functional homolog of human COL4A5 in nephrocytes, we carried out gene replacement experiments. The UAS-*COL4A5* transgenic fly line showed no changes in nephrocyte uptake capability (Figure 2; Supplemental Figure 1). However, when using this line to express human *COL4A5* in nephrocytes deficient for fly *Col4a1*, nephrocyte uptake of 10kD Dextran, FITC-albumin, and ANF-RFP (*Mhc*-ANF-RFP) significantly improved (Figure 2; Supplemental Figure 1). These findings suggest that in *Drosophila* nephrocytes Col4a1 is indeed the functional homolog of human COL4A5.

**Figure 2.**
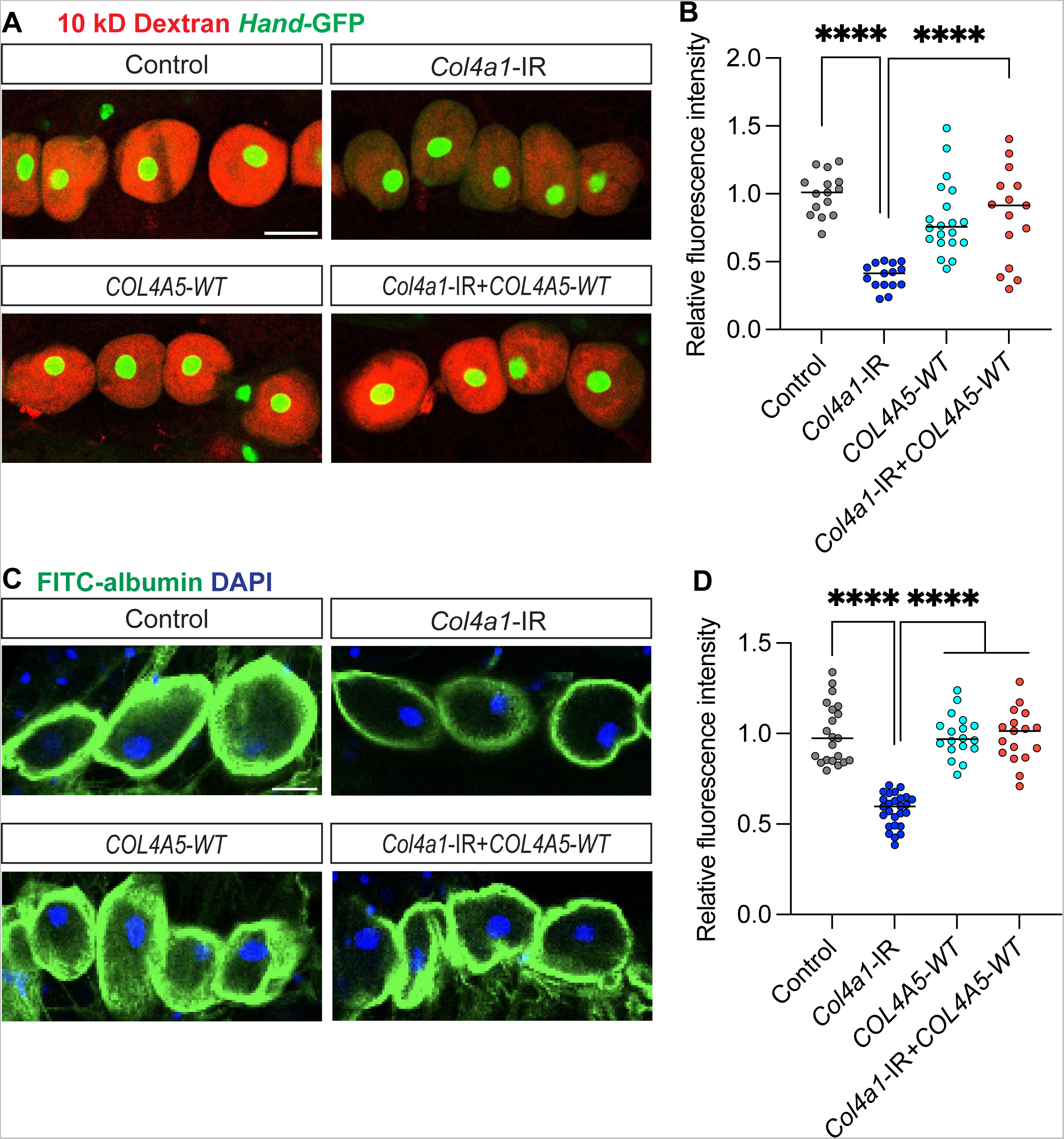
Human *COL4A5* can restore uptake function in *Col4a1* deficient nephrocytes. (**A**) Representative confocal images of 10 kD Dextran uptake (red) by female adult nephrocytes from Control, *Col4a1*-IR, *COL4A5-WT* (human reference *COL4A5*), and *Col4a1*-IR+*COL4A5-WT* fly lines. *Hand*-GFP transgene expression was visualized as green fluorescence concentrated in the nephrocyte nuclei. Scale bar: 50 µm. (**B**) Quantitation of the relative fluorescence intensity of 10 kD Dextran in (A). (**C**) Representative confocal images of FITC-albumin uptake by female adult nephrocytes from Control, *Col4a1*-IR, *COL4A5-WT* (human reference *COL4A5*), and *Col4a1*-IR+*COL4A5-WT* fly lines. DAPI (blue) indicates nuclei. Scale bar: 20 μm. (**D**) Quantitation of the relative fluorescence intensity of FITC-albumin in (C). (**A-D**) Flies: Control, (*Hand*-GFP/+; *Klf15*-Gal4/+); *Col4a1*-IR, (*Hand*-GFP/+; *Klf15*-Gal4/+; UAS-*Col4a1-*IR VDRC_28369/+); *COL4A5-WT* (*Hand*-GFP/+; *Klf15*-Gal4/UAS-*COL4A5-WT*); and *Col4a1*-IR+*COL4A5-WT*, (*Hand*-GFP/+; *Klf15*-Gal4/UAS-*COL4A5-WT*; UAS-*Col4a1-*IR VDRC_28369/+). (**B,D**) Statistical significance was defined as P<0.05 using one-way ANOVA with Tukey multiple comparisons test; (****) signifies P<0.0001.

### Nephrocyte *Col4a1* deficiency fly model to test pathogenicity of ClinVar *COL4A5* pathogenic variants associated with Alport syndrome

So far, we have shown that deficiency for fly C*ol4a1* causes nephrocyte defects and that the orthologous human *COL4A5* can ameliorate this phenotype, thus providing gene-level validation. For functional data to determine pathogenicity of patient variants associated with Alport syndrome we need variant-level validation. For this, we express human *COL4A5* alleles that carry a patient variant (*Klf15*-Gal4 driver for nephrocyte-specific expression) and assess if it can restore nephrocyte dysfunction induced by *Col4a1* deficiency (*Klf15*>RNAi) to the same extent as the human reference allele (wildtype).

First, three missense variants were identified in patients with Alport syndrome that have pathogenic classification in ClinVar, but without supporting in vitro or in vivo evidence: *COL4A5*-C1570S (NC1), *COL4A5*-L1655R (NC1), and *COL4A5*-G869R (triple-helix region) (Figure 3). Whereas expressing the human *COL4A5* reference allele (wildtype) restored nephrocyte uptake function, none of the three patient-variant *COL4A5* alleles could. For all three, the nephrocytes showed reduced ability to take up 10kD Dextran and FITC-albumin (Figure 4A-D). Transmission electron microscopy (TEM) showed structural differences in the nephrocyte basement membrane which was thick and irregular in *Col4a1*-IR flies, but which had structurally normal slit diaphragms (Figure 4E). These structural defects could be restored by expressing the human *COL4A5* reference allele (wildtype), but not by the patient variant *COL4A5* allele (*COL4A5*-C1570S) (Figure 4E).

**Figure 3.**
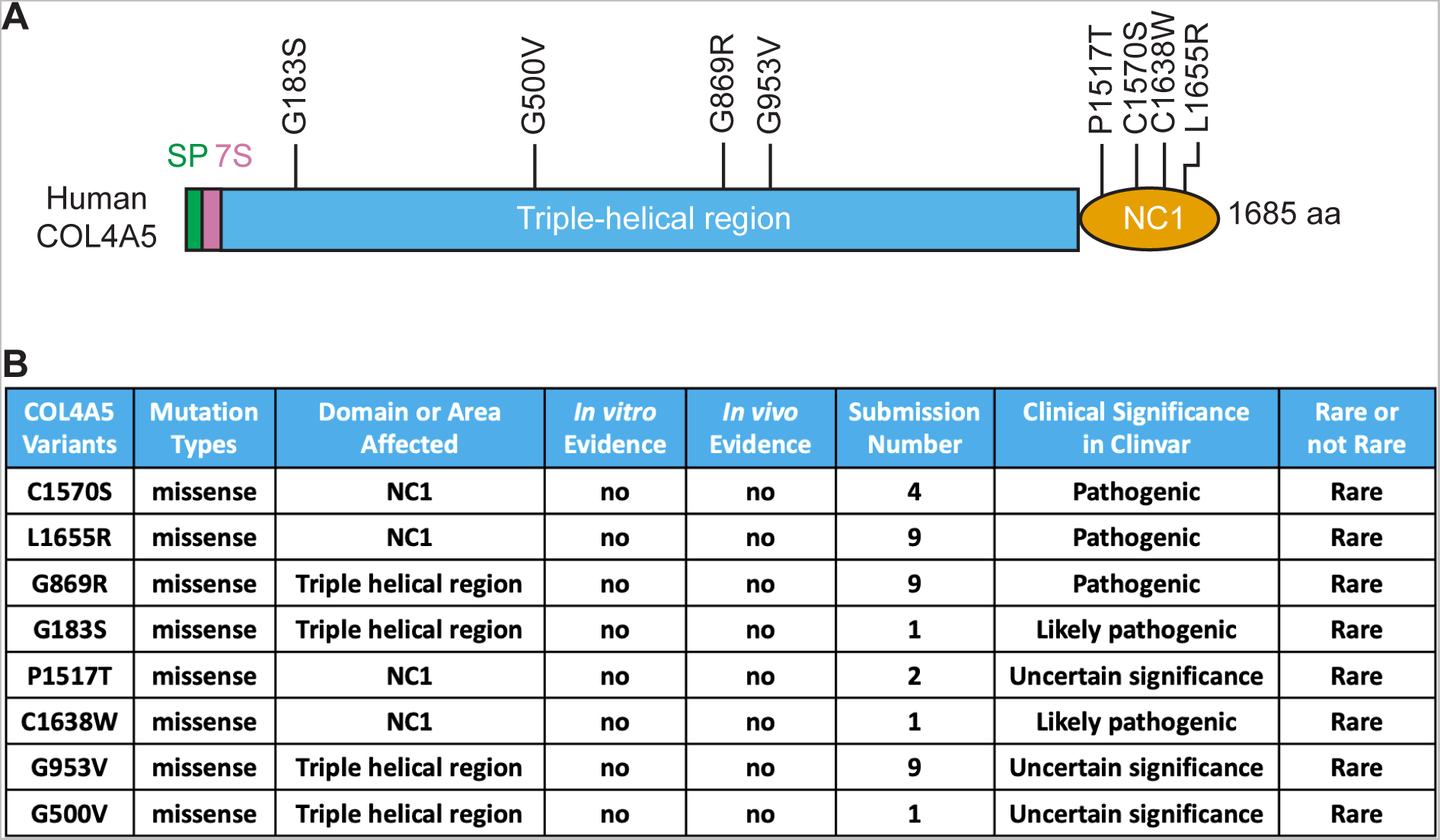
COL4A5 variants associated with Alport syndrome in patients (ClinVar) (**A**) Graphic representation of human COL4A5 with the location of the seven missense variants from ClinVar included in this study. 7S, seven Svedberg units; NC1, non-collagenous domain; SP, signal peptide. (**B**) A table with variant details obtained from ClinVar for the select human COL4A5 variants presented in (A).

**Figure 4.**
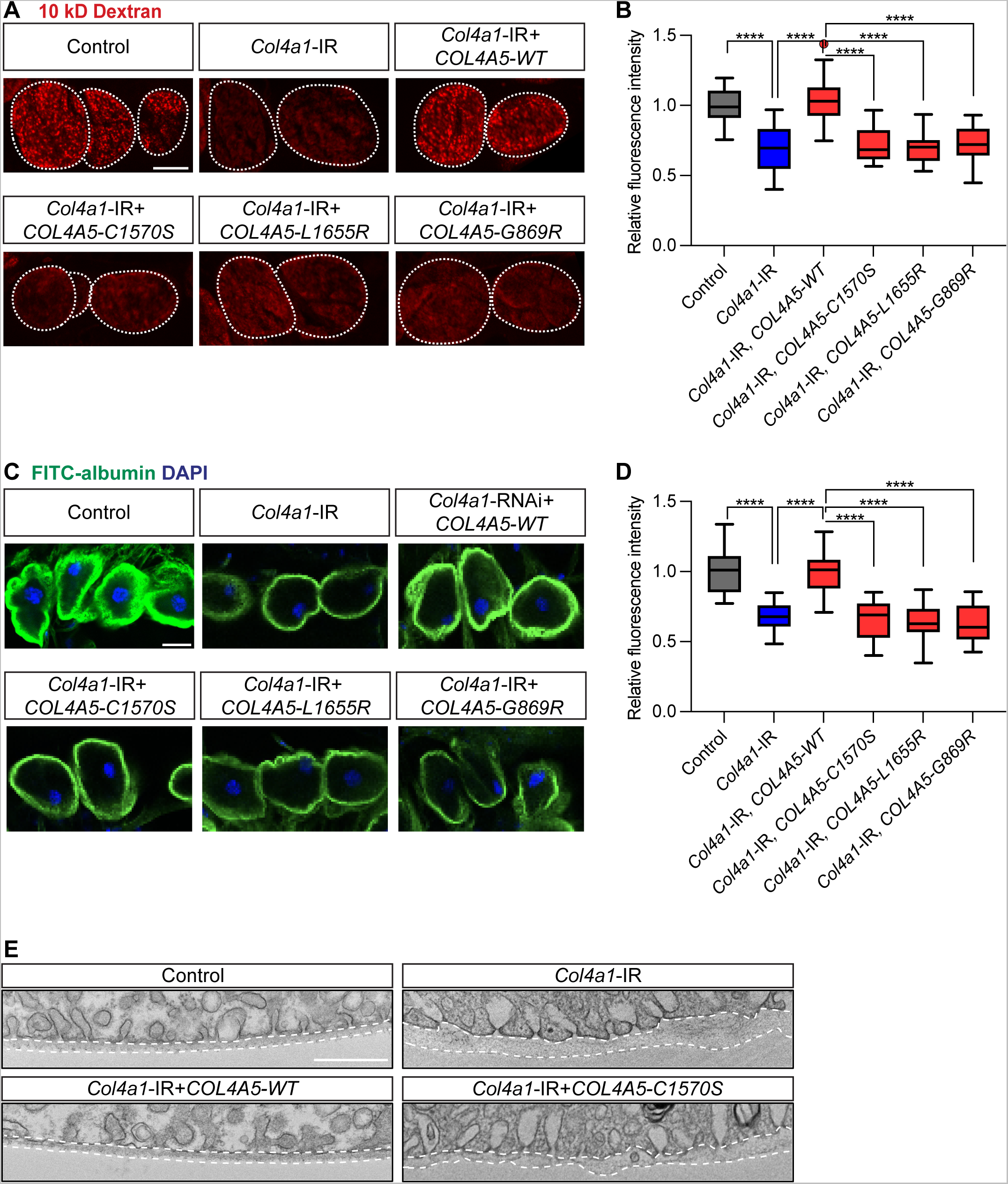
Pathogenic COL4A5 variants C1570S, L1655R, and G869R (ClinVar) could not restore uptake function in *Col4a1* deficient nephrocytes. (**A**) Representative confocal images of 10 kD Dextran uptake (red) by female adult nephrocytes from Control, *Col4a1*-IR, and *Col4a1*-IR+*COL4A5* (human reference, WT, or with patient variant) fly lines. Scale bar: 20 μm. (**B**) Quantitation of the relative fluorescence intensity of 10 kD Dextran in (A). (**C**) Representative confocal images of FITC-albumin uptake by female adult nephrocytes from Control, *Col4a1*-IR, and *Col4a1*-IR+*COL4A5* (human reference, WT, or with patient variant) fly lines. DAPI (blue) indicates nuclei. Scale bar: 20 μm. (**D**) Quantitation of the relative fluorescence intensity of FITC-albumin in (C). (**E**) Representative transmission electron microscopy (TEM) images of nephrocytes from Control, *Col4a1*-IR, and *Col4a1*-IR+*COL4A5* (human reference, WT, or with patient variant) fly lines. Scale bar: 500 nm. (**A-E**) Control, (*Klf15*-Gal4/+); *Col4a1*-IR, (*Klf15*-Gal4/+; UAS-*Col4a1-*IR VDRC_28369/+); *Col4a1*-IR+*COL4A5-WT*, (*Klf15*-Gal4/UAS-*COL4A5-WT*; UAS-*Col4a1*-IR VDRC_28369/+); *Col4a1*-IR+*COL4A5*-*C1570S*, (*Klf15*-Gal4/UAS-*COL4A5*-*C1570S* ; UAS-*Col4a1*-IR VDRC_28369/+); *Col4a1*-IR+*COL4A5*-*L1655R*, (*Klf15*-Gal4/UAS-*COL4A5*-*L1655R*; UAS-*Col4a1*-IR VDRC_28369/+); and *Col4a1*-IR+*COL4A5*-*G869R*, (*Klf15*-Gal4/UAS-*COL4A5*-*G869R*; UAS-*Col4a1*-IR VDRC_28369/+). (**B,D**) Statistical significance was defined as P<0.05 using one-way ANOVA with Tukey multiple comparisons test; (****) signifies P<0.0001; ns, not significant.

These results confirm pathogenicity of the variants, *i.e.*, all three showed an inability to restore normal nephrocyte function in *Col4a1*-deficient flies. The TEM findings suggest that these defects are due to structural issues in the nephrocyte basement membrane, reminiscent of patients with Alport syndrome in which the glomerular basement membrane (GBM) is irregular.

### Nephrocyte *Col4a1* deficiency fly model to test pathogenicity of *COL4A5* Alport syndrome variants classified as likely pathogenic or of uncertain significance

Based on the encouraging results from the *COL4A5* Alport syndrome variants with pathogenic classification in ClinVar (Figure 3; Figure 4), next we investigated missense variants with limited submissions or conflicting interpretations in ClinVar: *COL4A5*-G183S (triple-helix) and *COL4A5*-C1638W (NC1) classified as likely pathogenic variants; *COL4A5*-P1517T (NC1), *COL4A5*-G953V (triple-helix), and *COL4A5*-G500V (triple-helix) as variants of uncertain significance (Figure 3). Our variant-level assessment for nephrocyte function (10kD Dextran and FITC-albumin), showed that of the likely pathogenic *COL4A5* variants, G183S restored nephrocyte uptake function equal to *COL4A5*-WT, whereas C1638W function remained significantly reduced compared to *COL4A5*-WT expression in the *Col4a1*-IR flies (Figure 5). Among the three variants with uncertain significance, G1517T was unable to restore function in the *Col4a1* deficient fly nephrocytes indicating a pathogenic nature. However, *COL4A5*-G953V and G500V returned function back within *COL4A5*-WT levels (Figure 5), suggesting these variants are not pathogenic in kidney cells. Altogether, these data provide in vivo functional evidence to support the pathogenic nature of two variants and the benign nature of three variants in *COL4A5* with previously unresolved clinical significance in Alport syndrome.

**Figure 5.**
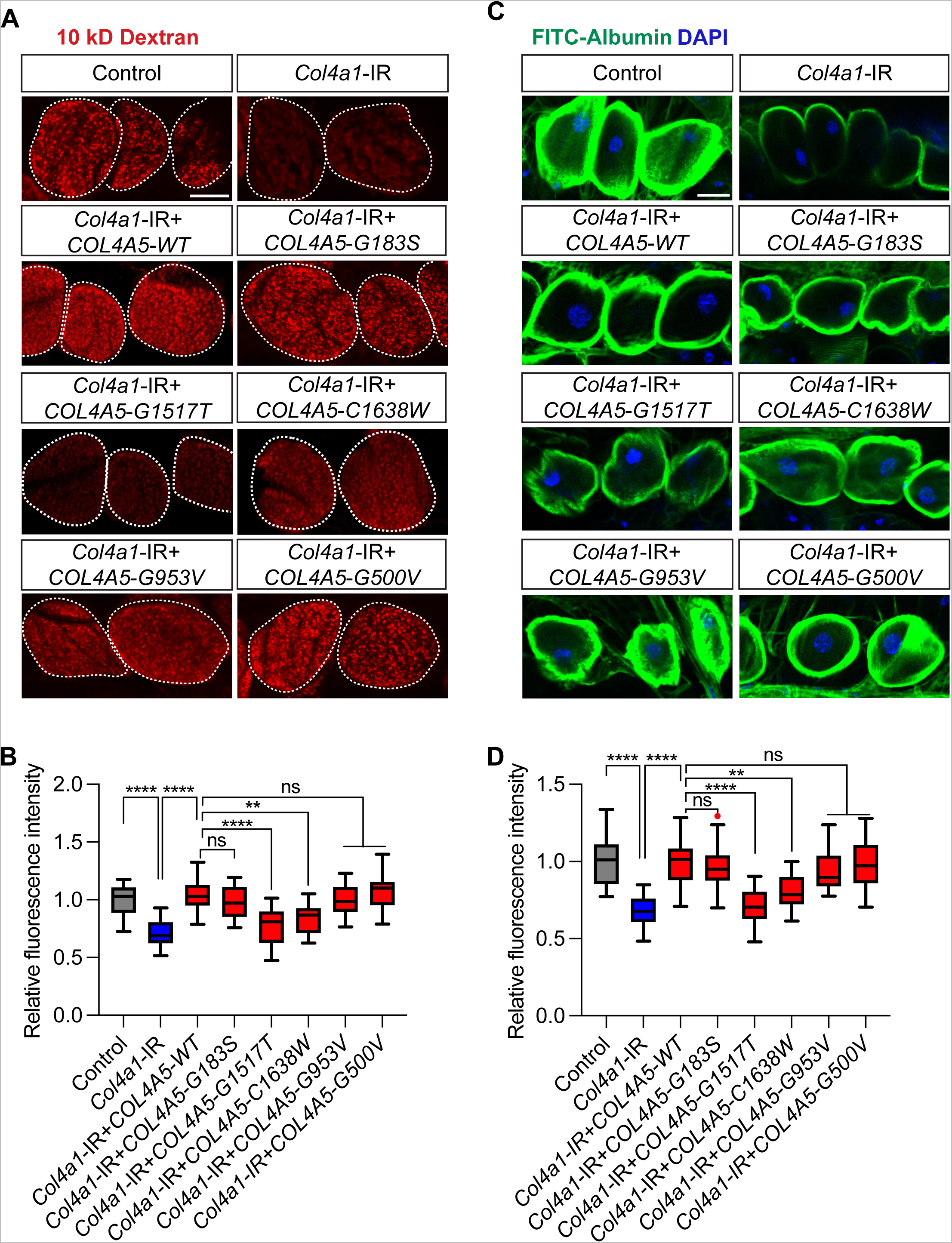
Assessment of likely pathogenic COL4A5 variants and those of uncertain significant associated with Alport syndrome (ClinVar) using the *Col4a1* deficient fly nephrocytes. (**A**) Representative confocal images of 10 kD Dextran uptake (red) by female adult nephrocytes from Control, *Col4a1*-IR, and *Col4a1*-IR + UAS-*COL4A5* (with patient variant) fly lines. Scale bar: 20 μm. (**B**) Quantitation of the relative fluorescence intensity of 10 kD Dextran in (A). (**C**) Representative confocal images of FITC-albumin uptake by female adult nephrocytes from Control, *Col4a1*-IR, and *Col4a1*-IR + UAS-*COL4A5* (with patient variant) fly lines. DAPI (blue) indicates nuclei. Scale bar: 20 μm. (**D**) Quantitation of the relative fluorescence intensity of FITC-albumin in (C). (**A-D**) Control, (*Klf15*-Gal4/+); *Col4a1*-IR, (*Klf15*-Gal4/+; UAS-*Col4a1-*IR VDRC_28369/+); *Col4a1*-IR+*COL4A5*-*G183S*, (*Klf15*-Gal4/UAS-*COL4A5*-*G183S*; UAS-*Col4a1-*IR VDRC_28369/+); *Col4a1*-IR+*COL4A5*-*G1517T*, (*Klf15*-Gal4/UAS-*COL4A5*-*G1517T*; UAS-*Col4a1-*IR VDRC_28369/+); *Col4a1*-IR+*COL4A5*-*C1638W*, (*Klf15*-Gal4/UAS-*COL4A5*-*C1638W*; UAS-*Col4a1-*IR VDRC_28369/+); and *Col4a1*-IR+*COL4A5*-*G953V*, (*Klf15*-Gal4/UAS-*COL4A5*-*G953V*; UAS-*Col4a1-*IR VDRC_28369/+). (**B,D**) Statistical significance was defined as P<0.05 using one-way ANOVA with Tukey multiple comparisons test; (***) signifies P<0.001, (****) signifies P<0.0001; ns, not significant.

### Nephrocyte *Col4a1* deficiency fly model to test pathogenicity of *COL4A5* Alport syndrome variants from the Chronic Kidney Disease in Children (CKiD) Study

ClinVar does not provide data on clinical presentation beyond the condition, *i.e.*, diagnosis. Therefore, we investigated two additional variants from the CKiD Study with clinical data available. The two participants in CKiD were diagnosed with Alport syndrome and both showed very fast progression. Patient 1 showed accelerated disease progression in late adolescence and patient 2 after 18 years of age. This rapid progression was captured by a decline in U25 estimated glomerular filtration rate (eGFR; ml/min|1.73m^2^), and an increase in urine protein:creatinine (UPCR; mg/mgCr) as a measure of kidney damage (Figure 6A). The CKiD study previously reported an average eGFR decline of 3.9% per year for those with nonglomerular (nearly all congenital) diagnoses (Pierce et al., 2011) and the linear decline for patient 1 and patient 2 was −13.9% and −5.2%, respectively. Both patients experienced a period with a sharp decline: The eGFR of patient 1 declined from 84 to 44 ml/min|1.73m^2^ from age 19 to 21; Patient 2 experienced a decline in eGFR from 92 to 71 ml/min|1.73m^2^ from age 17 to 19. Each patient carried a missense variant in the COL4A5 triple-helix domain: patient 1, *COL4A5*-G893A; patient 2, *COL4A5*-G1205D (Figure 6B). These were identified by targeted sequencing of 71 genes associated with nephrotic syndrome. These variants were classified as pathogenic in ClinVar based on independent patients (one patient per variant). We used our variant-level assessment in fly nephrocytes to provide functional data for these variant-disease associations. In *Col4a1*-deficient fly nephrocytes, the human COL4A5 carrying either patient variant (*COL4A5*-G893A, *COL4A5*-G1205D) was unable to restore the functional uptake deficit shown for 10 kD Dextran or the larger FITC-albumin particles (Figure 6C-F). Overall, these findings in fly reflect those in the patients with Alport syndrome and together provide in vivo functional support for the pathogenic nature of these identified CKiD patient variants.

**Figure 6.**
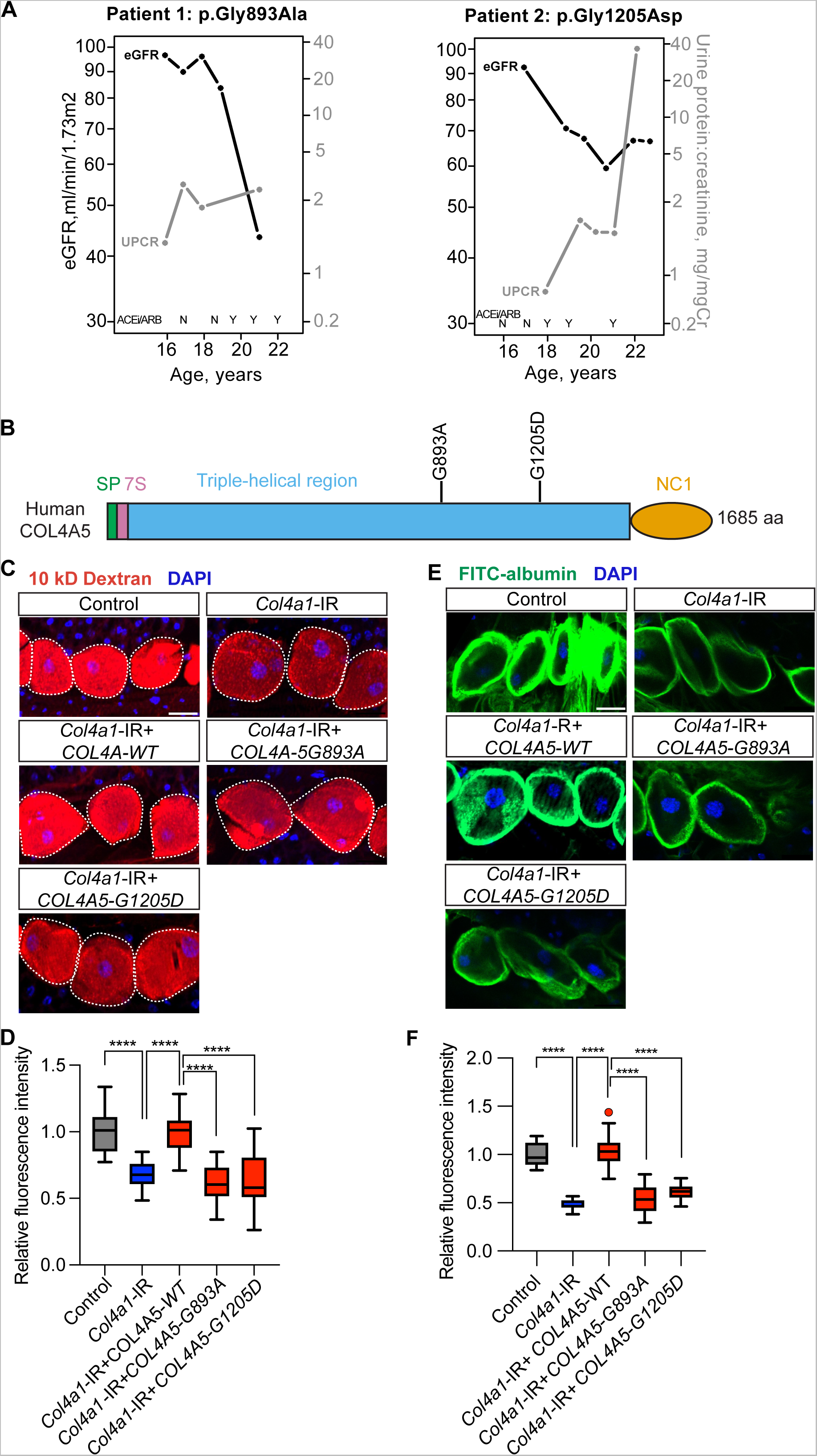
Assessment of CKiD patient variants in COL4A5 associated with Alport syndrome. (**A**) Longitudinal clinical estimated glomerular filtration rate (eGFR) and urine protein creatinine ratio (UPCR; Cr, creatinine) (time in years) for two patients who carry the new *COL4A5* variants (patient 1: p.Gly893Ala, G893A; patient 2: p.Gly1205Asp, G1205D). ACEi/ARB, self-reported presence (Y) or absence (N) of ACE inhibitor/ARB mediation use at time of measurement. (**B**) Graphic representation of human COL4A5 with the location of the two new patient variants; G893A and G1205D. (**C**) Representative confocal images of 10 kD Dextran uptake (red) by female adult nephrocytes from Control, *Col4a1*-IR, and *Col4a1*-IR + UAS-*COL4A5* (with patient variant) fly lines. DAPI (blue) indicates nuclei. Scale bar: 20 μm. (**D**) Quantitation of the relative fluorescence intensity of 10 kD Dextran in (C). (**E**) Representative confocal images of FITC-albumin uptake by female adult nephrocytes from Control, *Col4a1*-IR, and *Col4a1*-IR + UAS-*COL4A5* (with patient variant) fly lines. DAPI (blue) indicates nuclei. Scale bar: 20 μm. (**F**) Quantitation of the relative fluorescence intensity of FITC-albumin in (E). (**C-F**) Flies: Control, (*Klf15*-Gal4/+); *Col4a1*-IR, (*Klf15*-Gal4/+; UAS-*Col4a1-*IR VDRC_28369/+); *Col4a1*-IR+*COL4A5*-*G893A*, (*Klf15*-Gal4/UAS-*COL4A5*-*G893A*; UAS-*Col4a1-*IR VDRC_28369/+); and *Col4a1*-IR+*COL4A5*-*G1205D*, (*Klf15*-Gal4/UAS-*COL4A5*-*G1205D*; UAS-*Col4a1-*IR VDRC_28369/+). (**D,F**) Statistical significance was defined as P<0.05 using one-way ANOVA with Tukey multiple comparisons test; (*) signifies P<0.05, (****) signifies P<0.0001; ns, not significant.

## DISCUSSION

Collagen IV is an ancient protein that evolved over millions of years, yet its protein components are highly conserved (Fidler et al., 2017). Humans carry six genes that encode collagen IV proteins, whereas the typically leaner fly genome carries two, with vertebrate α1, α3, and α5 designated α1-like, and vertebrate α2, α4, and α6 designated α2-like (Fidler et al., 2017). Our data are in line with this designation. Fly nephrocytes deficient for *Col4a1* displayed irregular thickness of the basement membrane and significantly reduced uptake function (Figures 1-3; Supplementary Figure S1). These findings are reminiscent of the clinical observations in patients with Alport syndrome, in whom kidney biopsies have shown GBM thinning, thickening, and irregularities, with subsequent filtration defects, kidney dysfunction and ultimately failure (Barsotti et al., 2001; Kalluri et al., 1997; Longo et al., 2006; Noël, 2000). Moreover, when we expressed human reference (wildtype) *COL4A5* in these *Col4A1* deficient fly nephrocytes their phenotype resolved, and filtration function was restored (Figures 1-3; Supplementary Figure S1). Together the data indicate that in nephrocytes *Drosophila* Col4a1 is the functional homolog of human COL4A5 in podocytes and demonstrate that flies with *Col4a1*-deficient nephrocytes provide a relevant research model for Alport syndrome.

To functionally assess variants associated with Alport syndrome, we adapted our gene replacement *Drosophila* model for variant-level validation. Instead of human reference (wildtype) *COL4A5* in the *Col4a1*-deficient nephrocytes, we expressed human *COL4A5* that carried patient variants. We validated our approach by first assessing variants with strong pathogenic evidence, then we applied the method to seven additional variants of varying clinical significance (Figure 3; Figure 6). This provided in vivo functional evidence to support their pathogenic classification, corroborating the three pathogenic variants, and supporting classification or reclassification for the others (Figure 7). For two of the variants with limited prior information, we included longitudinal standardized clinical data for two independent patients. Both were diagnosed with Alport syndrome marked by rapid progression; the data in the fly supported pathogenicity for both variants (Figure 6). Altogether these data show that the *Drosophila* platform can provide a fast and economical screen to assess functional pathogenicity of variants in *COL4A5* associated with Alport syndrome. Knowing which variants are likely pathogenic, backed by functional data, can aid clinical diagnosis and help focus research efforts. Furthermore, our system could be readily adapted to include variants in *COL4A3* and *COL4A4*.

**Figure 7.**
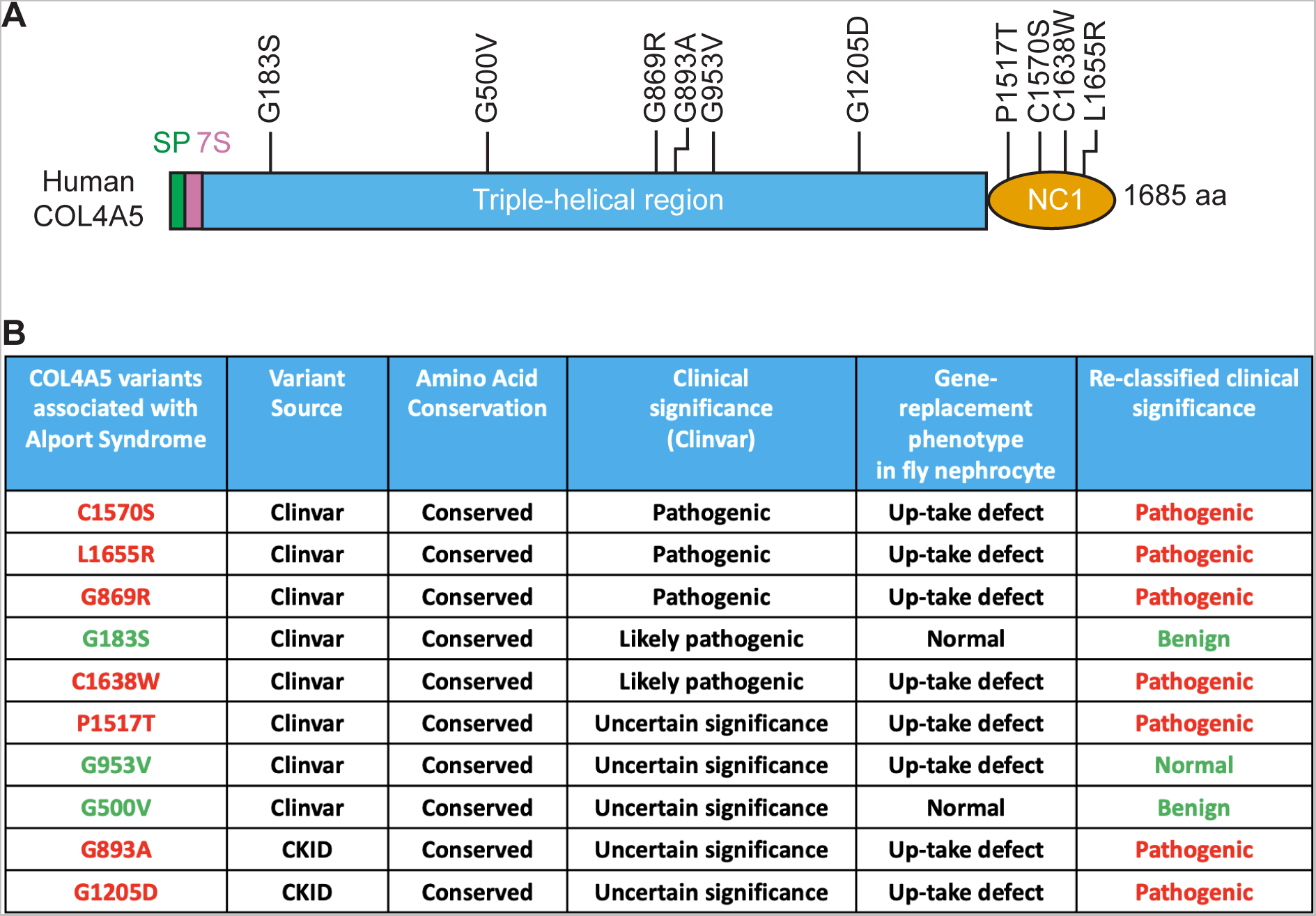
(re-)Classification of Alport syndrome associated *COL4A5* variants based on functional findings in *Drosophila* nephrocytes. (**A**) Graphic representation of human COL4A5 with the location of all the variants tested in this study. 7S, seven Svedberg units; NC1, non-collagenous domain; SP, signal peptide. (**B**) Summary data for the human COL4A5 variants (re-)classified based on fly in vivo 10 kD dextran and FTIC-albumin up take in *Drosophila* nephrocytes. Red font indicates pathogenic variant, green font indicates benign variant based on the current functional study in fly nephrocytes. CKiD, Chronic Kidney Disease in Children study; ClinVar, NCBI ClinVar.

The many variants in *COL4A5* and their varying pathogenic effect, put to question whether variant location in the type IV Collagen protein contributes to pathogenicity (Daga et al., 2022; Savige et al., 2021; Zhang et al., 2021). Using *Drosophila*, we tested variants that were either in the triple helix or in the NC1 (a.k.a., C4) domain, at which the three chains of the collagen fiber interact. All variants located in the NC1 domain were pathogenic, whereas some variants in the triple-helix maintained functionality, thus not supporting their pathogenic nature (Figures 4-6). Whereas Polymorphism Phenotyping (PolyPhen) prediction scores of the possible impact of an amino acid substitution on the structure and function of a human protein (Adzhubei et al., 2010) at times are in conflict with next-generation sequencing findings in Alport syndrome patient families (Artuso et al., 2012)—mammalian model systems (see review (Nikolaou and Deltas, 2022)) typically lack the large-scale screening capabilities needed to answer this question—*Drosophila* is well-suited for large-scale screens. Our fly system could be readily scaled to screen many variants across all COL4A5 protein domains. The findings could provide valuable insight into whether variants in certain domains are more detrimental than others, which could aid diagnostic application and the prioritization of research efforts.

The knowledge of genetic diagnosis in clinical management of kidney disease has been shown to improve patient outcomes (Groopman et al., 2019; Nestor et al., 2020). Particularly in light of phenocopies, for example, Alport syndrome may clinically present itself as focal segmental glomerulosclerosis (FSGS) or steroid-resistant nephrotic syndrome (SRNS) (Riedhammer et al., 2020). Different affected proteins might indicate a different pathomechanism, each of which requires a different targeted treatment approach. Aside from Alport syndrome in this study, flies have been successfully used to assess causality for genetic variants associated with diverse forms of kidney disease, including SRNS (example studies (Gonçalves et al., 2018; Hermle et al., 2017; Lovric et al., 2017; Milosavljevic et al., 2022; Odenthal et al., 2023; Zhang et al., 2013; Zhao et al., 2019; Zhu et al., 2017)). Therefore, our *Drosophila* in vivo nephrocyte functional screening system for patient derived genetic variants could be applied to clinical variants associated with other nephropathies.

## MATERIALS AND METHODS

### *Drosophila* stocks and maintenance

All fly stocks were maintained at 25°C with 12 h light-dark cycles and 60% humidity, on a standard diet (Meidi Laboratories, MD). The *Drosophila UAS-Col4a1-*IR lines were obtained from the Bloomington Drosophila Stock Center (Bloomington, IN) (BDSC ID: 44520) and the Vienna Drosophila Resource Center (Vienna, Austria) (VDRC ID: 28369). The following *Drosophila* lines with prior publications have been used: *Hand*-GFP (Han and Olson, 2005), +/CyO-*Dfd*-EYFP (Le et al., 2006), and *Mhc*-ANF-RFP (Zhang et al., 2013).

### Generation *Drosophila Klf15*-Gal4

To generate the *Klf15*-Gal4 transgenic line, a 2.1 kb *Klf15* promoter region was PCR amplified using the following primers (Klf15F 5’-3’: ATCTGTTAACGAATTCGTCCTCGGATTTGCTTCGTAAATACTTGC and Klf15R 5’-3’: TCTTTTCGCCGGATCCGATCGCAAATGAGCGGACTCCAGTC) and cloned into the pPTGAL vector between EcoRI and BamHI restriction sites. The plasmid was sequence verified. Microinjection was performed by Rainbow Transgenic Flies (CA, USA).

### Generation of transgenic *Drosophila* carrying *COL4A5* wildtype and select variants

The following *Drosophila* lines were generated in house to carry Alport syndrome associated variants in human *COL4A5*: UAS-*COL4A5*-*C1570S*, UAS-*COL4A5*-*L1655R*, UAS-*COL4A5*-*G869R*, UAS-*COL4A5*-*G183S*, UAS-*COL4A5*-*G1517T*, UAS-*COL4A5*-*C1638W*, UAS-*COL4A5*-*G953V*, UAS-*COL4A5*-*G500V*, UAS-*COL4A5*-*G893A*, and UAS-*COL4A5*-*G1205D*. The cDNA corresponding to human wildtype *COL4A5* (GenBank accession no. NM_033380.3) was obtained commercially (genomics-online.com, ABIN4071174), subcloned into the pUASt-attB vector, then sequenced to ensure sufficient quality. Next, oligonucleotide primers were designed to introduce the respective mutant sites using the PCR-based In-Fusion cloning technique (Takara Bio, Japan). The transgenes with human reference wildtype *COL4A5* and select patient-derived variants were introduced to the 51C attP landing site on the second chromosome by Rainbow Transgenic Flies (CA, USA). These flies were then crossed with +/CyO-*Dfd*-EYFP flies to balance against yellow fluorescence in the head (*Dfd*-EYFP) at the embryonic and larval fly stages and curly wing (CyO) in the adult flies.

### Dextran uptake assay

Flies carrying *Hand-GFP* and *Klf15-Gal4* transgenes were crossed with flies carrying the *UAS*-RNAi transgenes at 25°C. Dextran uptake was assessed in adult flies, one-day post-emergence, by dissection of the pericardial nephrocytes in Schneider’s Drosophila Medium (Thermo Fisher, 21720024) and examination of the cells by fluorescence microscopy after a 20 min incubation with Texas Red labeled dextran (10 kD, 0.02 mg/ml; Thermo Fisher, D1828) (Wang et al., 2021). For dextran absorption, female adults were dissected in Schneider’s Drosophila Medium and incubated in the dextran solution for 30 min at room temperature. Then the samples were fixed in 4% PFA in phosphate buffered saline (1xPBS) for 30 min, followed by a wash with 0.2% Triton x-100 in 1xPBS (1xPBST), *Hand*-GFP or DAPI were used to visualize the nephrocyte nuclei (DAPI: 10 min incubation in DAPI solution, 0.5 mg/ml; Thermo Fisher, D1306), followed by two washes with 1xPBST, then once with 1xPBS, for 10 min each.

### FITC-albumin uptake assay

For FITC-albumin absorption, 1-day-old female adults were dissected in Schneider’s Drosophila medium (Thermo Fisher, #21720024) and incubated in FITC-albumin solution (10 mM; Sigma, A9771) for 1 min at room temperature. Then the samples were fixed in 4% PFA in phosphate buffered saline (1xPBS) for 30 min, followed by a wash with 0.2% Triton x-100 in 1xPBS (1xPBST), a 10 min incubation in DAPI solution (0.5 mg/ml; Thermo Fisher, D1306), followed by two washes with 1xPBST, then once with 1xPBS, for 10 min each.

### *Mhc*-ANF-RFP

*Mhc-ANF-RFP; Klf15*-Gal4 virgins were crossed with *w^1118^*or *Col4a1*-IR (RNA interference) and UAS-*COL4A5 Drosophila* lines. Around 8 h following eclosion, the female adults were dissected in Schneider’s Drosophila Medium (Thermo Fisher, 21720024), then fixed with 4% PFA in 1xPBS for 30 min, followed by a wash with 1xPBS, a 10 min incubation in DAPI solution (0.5 mg/ml; Thermo Fisher, D1306), followed by two washes with 1xPBST, then once with 1xPBS, for 10 min each. Samples were then mounted using Vectashield (Vector Laboratories, H-1000-10) mounting medium.

### Confocal microscopy

Confocal imaging was performing using a ZEISS LSM 900 microscope using a 20X objective and ZEISS Zen 3.0 (Blue edition) acquisition software. For quantitative comparison of intensities, settings were chosen to avoid oversaturation (by limiting the oversaturated pixels visualized using Range Indicator in Zen Blue) and applied across image for all the samples within an assay. Images were processed using ImageJ software (version 1.53t; Fiji version 2.9.0) (Schneider et al., 2012).

### Transmission electron microscopy (TEM)

TEM was carried out using standard procedures. Briefly, one-day-old adult flies of the indicated genotypes were dissected in artificial hemolymph and fixed in 8% paraformaldehyde for 10 min. Then the tissues were further trimmed in 1xPBS. The trimmed samples were transferred into fixation buffer containing 4% paraformaldehyde and 2.5% glutaraldehyde. The samples were further processed and analyzed using a FEI Tecnai T12 TEM (Wang et al., 2021) at the Electron Microscopy Core Imaging Facility at the Center for Innovative Biomedical Resources (CIBR) (University of Maryland School of Medicine, MD, USA).

### Statistical analyses

Prism9 (GraphPad; version 9.5.1) was used to perform the statistical analysis and graphical display of the data. All experiments were repeated at least three times. One-way ANOVA with Tukey’s multiple comparisons test was used to determine statistical significance. The difference between two groups was defined as statistically significant for the following p values: *<0.05, **<0.01, ***<0.001, ****<0.0001.

### Patients

The Chronic Kidney Disease in Children (CKiD) study is an ongoing prospective longitudinal multicenter observational cohort of children with a previous diagnosis of kidney disease and mild to moderate CKD. Participants attended annual visits to contribute biological samples and answer questionnaires regarding general health and therapy use (a full description of the study design has been published (Furth et al., 2006)). All participants and families provided informed consent/assent, and the study protocols were approved by local institutional review boards. The CKiD study is carried out conform the Declaration of Helsinki. Patients with diagnoses of proteinuric kidney diseases were chosen for targeted sequencing of 71 genes associated with nephrotic syndrome. A filtering pipeline was applied to the variants called to identify participants with a putative Mendelian form of nephrotic syndrome. Qualifying missense variants were those with (1) a sample allele frequency less than 0.4%, (2) maximum allele frequency in gnomAD (Chen et al., 2022) less than 0.1% or missing, (3) at least 2 out of 3 functional prediction programs (MutationTaster, Polyphen, SIFT) predicted as damaging, (4) Genomic Evolutionary Rate Profiling (GERP) score more than 4, and (5) sequence read counts more than 20. Qualifying loss-of-function variants were those with (1) sample allele frequency less than 1%, (2) maximum allele frequency in gnomAD less than 0.1% or missing, and (3) sequence read counts more than 20. Variants that met this threshold were then evaluated for their ACMG classification of pathogenicity using Varsome (Kopanos et al., 2019).

### eGFR, UPCR, and ACEi/ARB

To characterize disease progression for each participant, repeated measures of serum creatinine and cystatin c-based U25 eGFR (ml/min|1.73m^2^) (Pierce et al., 2011) and urine protein:creatinine ratio (mg/mg creatinine) were plotted on the log scale, using age as the time scale along with self-reported ACEi/ARB therapy use at each measurement.

## ACKNOWLEDGMENTS

We thank the children and families of CKiD for their participation, time, and commitment. Data in this manuscript were collected by the Chronic Kidney Disease in Children prospective cohort study (CKiD) with clinical coordinating centers (Principal Investigators) at Children’s Mercy Hospital and the University of Missouri – Kansas City (Bradley Warady, MD) and Children’s Hospital of Philadelphia (Susan Furth, MD, PhD), Central Biochemistry Laboratory (George Schwartz, MD) at the University of Rochester Medical Center, and data coordinating center (Alvaro Muñoz, PhD and Derek Ng, PhD) at the Johns Hopkins Bloomberg School of Public Health. The CKiD Study is supported by grants from the National Institute of Diabetes and Digestive and Kidney Diseases, with additional funding from the Eunice Kennedy Shriver National Institute of Child Health and Human Development, and the National Heart, Lung, and Blood Institute (U01 DK066143, U01 DK066174, U24 DK082194, U24 DK066116). The CKiD website is located at https://statepi.jhsph.edu/ckid and a list of CKiD collaborators can be found at https://statepi.jhsph.edu/ckid/site-investigators/. We also thank the Bloomington Drosophila Stock Center (BDSC) based at Indiana University and the Vienna Drosophila Resource Center (VDRC) part of the Vienna BioCenter Core Facilities for the *Drosophila* stocks. We extend our thanks to the Electron Microscopy Core Imaging Facility at the Center for Innovative Biomedical Resources (CIBR) (University of Maryland School of Medicine, MD, USA) for their support in TEM image acquisition.

## DATA AVAILABILITY

All relevant data can be found within the article and its supplementary information. The materials that support the findings of this study are available from the corresponding author upon reasonable request. Due to multiple participating sites, please contact Dr. Sampson for information about genetic sequence data (CKiD) and he will work with you to share what is possible under the consent.

## AUTHOR CONTIBUTIONS

P.W. and Z.H. designed the study; J.D., P.W., and Y.Z. carried out the experiments; J.D., P.W., Y.Z., J.vdL., and Z.H. analyzed and/or interpreted the data; J.L.Y. and D.F. carried out patient sequencing for CKiD; B.A.W., S.L.F., D.K.N. and M.G.S. collected, analyzed and interpreted the clinical data from CKiD; J.D., P.W., Y.Z., and D.K.N. prepared the figures; J.vdL., and Z.H. drafted and revised the manuscript; the manuscript has been critically reviewed and the final version approved by all authors.

## SUPPLEMENTARY FIGURE LEGENDS

**Supplementary Figure S1. Human *COL4A5* can restore ANF-RFP uptake in *Col4a1* deficient nephrocytes** (**A**) Representative confocal images of ANF-RFP uptake (red) by female adult nephrocytes from Control, *Col4a1*-IR, UAS-COL4A5, and *Col4a1*-IR; *COL4A5-WT* (human reference) *Mhc*-ANF-RFP fly lines. DAPI (blue) indicates nuclei. *Mhc*-ANF-RFP, *Myosin heavy chain* (*Mhc*) promoter region drives expression of full-length Natriuretic peptide A (Rnor\Nppa) cDNA tagged with the DsRed(T4) fluorescent protein. Scale bar: 50 μm. (**B**) Quantitation of the relative fluorescence intensity of ANF-RFP in (A). Statistical significance was defined as P<0.05 using one-way ANOVA with Tukey multiple comparisons test (****) signifies P<0.0001 (**A,B**) Flies: Control, (*Hand*-GFP, *Mhc*-ANF-RFP/+; *Klf15*-Gal4/+); *Col4a1*-IR, (*Hand*-GFP, *Mhc*-ANF-RFP/+; *Klf15*-Gal4/+; UAS-*Col4a1*-IR VDRC_28369/+); *COL4A5-WT* (*Hand*-GFP, *Mhc*-ANF-RFP/+; *Klf15*-Gal4/UAS-*COL4A5-WT*); and *Col4a1*-IR+*COL4A5-WT*, (*Hand*-GFP, Mhc-ANF-RFP/+; *Klf15*-Gal4/UAS-*COL4A5-WT*; UAS-*Col4a1*-IR VDRC_28369/+).

